# B LYMPHOCYTES, BUT NOT DENDRITIC CELLS, EFFICIENTLY HIV-1 *TRANS*-INFECT NAÏVE CD4^+^ T CELLS: IMPLICATIONS FOR THE VIRAL RESERVOIR

**DOI:** 10.1101/2020.10.22.351627

**Authors:** Abigail Gerberick, Diana C. DeLucia, Paolo Piazza, Mounia Alaoui-El-Azher, Charles R. Rinaldo, Nicolas Sluis-Cremer, Giovanna Rappocciolo

## Abstract

Insight into the establishment and maintenance of HIV-1 infection in resting CD4^+^ T cell subsets is critical for the development of therapeutics targeting the HIV-1 reservoir. Although the frequency of HIV-1 infection, as quantified by the frequency of HIV-1 DNA, is lower in CD4^+^ naïve T cells (T_N_) compared to the memory T cell subsets, recent studies have shown that T_N_ cells harbor a large pool of replication-competent virus. Interestingly, however, T_N_ cells are highly resistant to direct (*cis*) HIV-1 infection *in vitro*, in particular to R5-tropic HIV-1, as T_N_ cells do not express CCR5. In this study, we investigated whether T_N_ cells could be efficiently HIV-1 *trans*-infected by professional antigen-presenting B lymphocytes and myeloid dendritic cells (DC) in the absence of global T cell activation. We found that B cells, but not DC, have a unique ability to efficiently *trans* infect T_N_ cells *in vitro*. In contrast, both B cells and DC mediated HIV-1 *trans* infection of memory and activated CD4^+^ T cells. Moreover, we found that T_N_ isolated from HIV-1-infected nonprogressors (NP) harbor significantly disproportionately lower levels of HIV-1 DNA compared to T_N_ isolated from progressors. This is consistent with our previous finding that APC derived from NP do not efficiently *trans*-infect CD4^+^ T cells due to alterations in APC cholesterol metabolism and cell membrane lipid raft organization. These findings support that B cell-mediated *trans* infection of T_N_ cells with HIV-1 has a more profound role than previously considered in establishing the viral reservoir and control of HIV-1 disease progression.

**Importance:** The latent human immunodeficiency virus type 1 (HIV-1) reservoir in persons on antiretroviral therapy represents a major barrier to a cure. Although most studies have focused on the HIV-1 reservoir in the memory T cell subset, replication competent HIV-1 has been isolated from naïve T cells, and CCR5-tropic HIV-1 has been recovered from CCR5^neg^T_N_ cells from ART-suppressed HIV-1-infected individuals. In this study, we showed that CCR5^neg^T_N_ cells are efficiently *trans* infected with R-5 tropic HIV-1 by B lymphocytes, but not by myeloid dendritic cells. Furthermore, we found that T_N_ isolated from NP harbor no or significantly less copies of HIV-1 DNA compared to ART-suppressed progressors. These findings support that B cell-mediated *trans* infection of T_N_ cells with HIV-1 has a more profound role than previously considered in establishing the viral reservoir and control of HIV-1 disease progression. Understanding the establishment and maintenance of the HIV-1 latent reservoir is fundamental for the design of effective treatments for viral eradication.

## Introduction

Latently infected resting CD4^+^ T cells constitute a major reservoir of persistent HIV-1 infection. Strategies that lead to a significant reduction or elimination of this reservoir could help in the development of either a functional or sterilizing cure (1–4). The CD4^+^ T cell population is heterogenous, broadly comprised of naïve (T_N_) and memory cells that differ in lifespan, proliferative capacity, localization and HIV-1 coreceptor expression. Memory cells are further categorized by various stages of differentiation, namely central memory (T_CM_), transitional memory and effector memory. The latent HIV-1 reservoir in memory T cell subsets has been extensively studied, whereas T_N_ cells have been largely overlooked (5, 6). Although resting T_N_ cells are highly resistant to direct, *cis* infection with HIV-1 *in vitro*, we and others have shown that HIV-1 DNA is detectable in T_N_ of viremic and virus-suppressed individuals (7, 8). While the frequency of HIV-1 infection in T_N_ is lower compared to T_CM_ cells, as much or more virus is produced by T_N_ cells after reactivation with latency reversing agents (LRAs) (9). Moreover, paradoxically, CCR5-tropic HIV-1 has been recovered from T_N_ despite the fact that they do not express the CCR5 coreceptor (9–12).

HIV-1 can infect its target cells via direct, *cis* infection or through a cell-to-cell transfer which can result in *trans* infection (13–15). This latter mechanism has been extensively described as mediated by professional antigen-presenting cells (APC), i.e., monocytes/macrophages, myeloid dendritic cells (DC) and B lymphocytes. Indeed, HIV-1 *trans* infection mediated by APC is 10- to 1000-fold more efficient than passive, *cis* dissemination of virions through the extracellular milieu (16, 17). We have previously shown that APC derived from HIV-1-infected nonprogressors (NP) do not efficiently transfer HIV-1 to CD4^+^ T cells due to alterations in APC cholesterol metabolism and cell membrane lipid raft organization (18, 19). In the present study, we show that B lymphocytes, but not DCs, have the exclusive ability to efficiently *trans* infect T_N_ cells with CCR-5 tropic HIV-1. Furthermore, T_N_ isolated from HIV-1 NP harbor significantly lower levels of HIV-1 DNA compared to T_N_ isolated from HIV-1 progressors (PR). These findings support that B cell-mediated *trans* infection of T_N_ cells with HIV-1 has a more profound role than previously considered in establishing the viral reservoir and control of HIV-1 disease progression.

## Materials and Methods

### Ethics Statement

Biological samples were acquired and studied from consented individuals according to University of Pittsburgh International Review Board approved protocols. All recruited participants were over the age of 18 and provided written consent prior to sample collection or use.

### Cohort

Experiments were performed using peripheral blood mononuclear cells (PBMC) obtained from Pittsburgh Blood Bank anonymous donors (HIV-1 negative, N=6) or archived PBMC obtained from 7 HIV-1 infected NP and 7 HIV-1 infected PR enrolled in the Pittsburgh portion of the MACS/WIHS Combined Cohort Study (MWCCS). The NP cohort consisted of 3 long term NP (LTNP, CD4^+^ T cells counts >500c/mm^3^ over >7 years post infection), 3 elite controllers (EC, undetectable viral load >7 years post infection) and 1 viremic controller (VC, at least two viral load measures below 2000 copies HIV-1 RNA/ml)).

### Generation of CD4^+^ T cell subsets

Naïve and central memory CD4^+^ T lymphocytes were selected from resting PBMC by magnetic bead negative selection according to the manufacturer’s instructions (Milenyi Biotech). T_N_ CD4^+^ T cells were defined as CD45RA^+^ CCR7^+^ CCR5^-^, while T_CM_ CD4^+^ T cells were defined as CD45RA^-^ CCR7^+^ CCR5^+^. The relative purity of the separated fractions was determined by flow cytometry.

### Cell isolation and culture

CD4^+^ T lymphocytes, B lymphocytes, and CD14^+^ monocytes were positively selected from PBMC using anti-CD4, -CD19, or -CD14 monoclonal antibody (MAb)- coated magnetic beads (Miltenyi Biotech). Immature DC (iDC) were derived from CD14^+^ monocytes cultured with 1,000 U/ml granulocyte-macrophage colony-stimulating factor (GM-CSF, Miltenyi Biotech) and 1,000 U/ml recombinant human interleukin-4 (rhIL-4) for 5 days in AIM-V medium, with additional GM-CSF and rhIL-4 on day 3. Mature DC (mDC) were derived from iDC by addition of 0.1 μg/ml trimeric CD40L (Enzo) on day 5 and cultured for an additional 2 days. Prior to coculture, CD4^+^ T cells and B cells were activated for 48h with 10U/ml IL-2 (Roche) and 2 μg/ml phytohemagglutinin (PHA, Sigma) or 1,000 U/ml rhIL-4 and 0.1 μg/ml trimeric CD40L (Enzo), respectively. CD4+ T_N_ or T_CM_ cells were treated with either 100nM CCL-19 (R&D Systems) or 10U/ml IL-2 and 2 μg/ml PHA as described. (12, 15)

### Cell phenotyping

Cells were assessed for surface protein expression by flow cytometry. B cell + T_N_ and DC + T_N_ cocultures or T_N_ cells alone were incubated with LIVE/DEAD fixable aqua viability cell stain kit (Invitrogen) for 20 min and then subsequently incubated with monoclonal antibodies (mAb) against CD3 (APC-H7), CD4 (V450), CCR5 (PE), CD45RA (PE-CF594), CCR7(APC), and CD27 (FITC) for 20 min. Cells were fixed with 1% paraformaldehyde (PFA), acquired with a BD LSR Fortessa and analyzed with FlowJo V10. The gating strategy is described in Supplemental Figure 1. Mature DC were also stained for siglec-1 expression (CD169-PE)

Virus stock titration and experimental p24 measurements were acquired by enzyme-linked immunosorbent assay (ELISA) using the HIV-1 p24 antigen capture immunoassay (Leidos Biomedical Research, Frederick National Laboratory for Cancer Research) per the manufacturer’s instructions. HIV-1 Gag p24 was also evaluated in *trans* and *cis* infection cocultures by flow cytometry. Briefly, cocultures were harvested and incubated with LIVE/DEAD fixable aqua viability stain kit (Invitrogen) for 20 min and then incubated for surface staining with monoclonal antibodies against CD3 (APC-H7), CD4(PE), CD19(PE-CF594) for 20 min. Cells were then permeabilized with PermII buffer (BD) for 20 min, washed and then incubated with anti-HIV-1 p24 antibody Kc57-FITC (Coulter), incubated for 20 min, washed and resuspended in 1% PFA prior to analysis with a BD LSR Fortessa. Acquired data were analyzed with FlowJo V10. The gating strategy is described in supplemental figure 2

### *Trans* and *cis* infection

R5-tropic HIV-1_BaL_, grown in and purified from PM1 cells (20) (American Type Culture Collection) was used for *cis* and *trans* infection experiments. A patient isolate R5-tropic-HIV-1 BX08(92FR_BX08) used in *trans* and *cis* experiments was obtained from the NIH AIDS Reagents Program, Division of AIDS, NIAID, NIH:HIV-1 BX08(92FR_BX08) virus(cat#11420) from Dr. Victoria Polonis (21). (i) *Trans infection*: 1 x 10^6^ APC were incubated with a low concentration of HIV-1_BaL_ or HIV-1BX08 (m.o.i. 10^−3^) for 2h at 37°C and then washed 3 times with cold medium. Virus-loaded APC were cocultured with autologous CD4^+^ T cell targets at 1:10 effector/target ratio in R10 medium. (ii) *Cis infection*: 1 x 10^6^ activated CD4+ T cells were incubated with a high (10^−1^) m.o.i. of HIV-1^BaL^ or HIV-1BX08 and cultured independently. HIV-1 Gag p24 levels were quantified in cell-free supernatants at days 4, 8, and 12 post coculture. In some experiments, stimulated B cells were incubated with 20μg/ml anti–DC-SIGN mAb (clone 120507, R&D system) or mouse IgG (R & D Systems) for 30 min at 4°C prior to incubation with virus. In some experiments, T_N_ were incubated with maraviroc (1μM) as previously described (22)

### Reactivation of latent HIV-1 from APC-TN cocultures

Eight days after the start of APC-T_N_ cocultures, cells were treated with 10 nM phorbol myristate acetate (PMA; Sigma-Aldrich) and 10 μg/ml PHA (PMA-PHA). Supernatants were collected at day 11, 14 and 17. Levels of HIV-1Gag p24 were then tested by ELISA. Parallel untreated cultures were used as control.

### Quantification of total HIV-1 DNA

Total HIV-1 DNA in CD4^+^ T cells was quantified as described previously (23).

### Statistics

Data were analyzed by one-way analysis of variance. Student t tests were used to compare two groups. GraphPad prism 7.0 Software was used for statistical analysis.

## Results

### B cells *trans* infect CD4^+^ T cells with high efficiency

B cells activated with CD40L and IL4, which mimics signals received from activated CD4^+^ T cells, express the C-type lectin DC-specific intercellular adhesion molecule-3-grabbing non-integrin (DC-SIGN) and can capture HIV-1, leading to *trans* infection of CD4^+^ T cells (24). In this study, we first extended this finding by demonstrating that B cells or DC loaded with a low, 10^−3^ m.o.i. of R5-tropic HIV-1_BaL_ could *trans* infect PHA/IL2 activated CD4^+^ T cells, with the efficiency of *trans* infection being significantly greater for B cells compared to DC (Figure 1A). This is of importance because, as we showed previously, only about 10-15% of activated B cells express DC-SIGN, compared to 100% of DC, therefore making B cells extraordinarily efficient in mediating HIV-1 *trans* infection. In agreement with our previous findings (15), CD4^+^ T cells were refractory to *cis* HIV-1 infection at the same low 10^−3^ m.o.i., but were productively infected with a 100-fold greater dose of 10^−1^ m.o.i. (Figure 1B). We conclude from these data that B lymphocytes, activated by two surrogates for CD4^+^ T helper cells, i.e., IL4 and CD40L, are more efficient than myeloid DC in mediating HIV-1 *trans* infection of activated CD4^+^ T lymphocytes.

**Figure 1.**
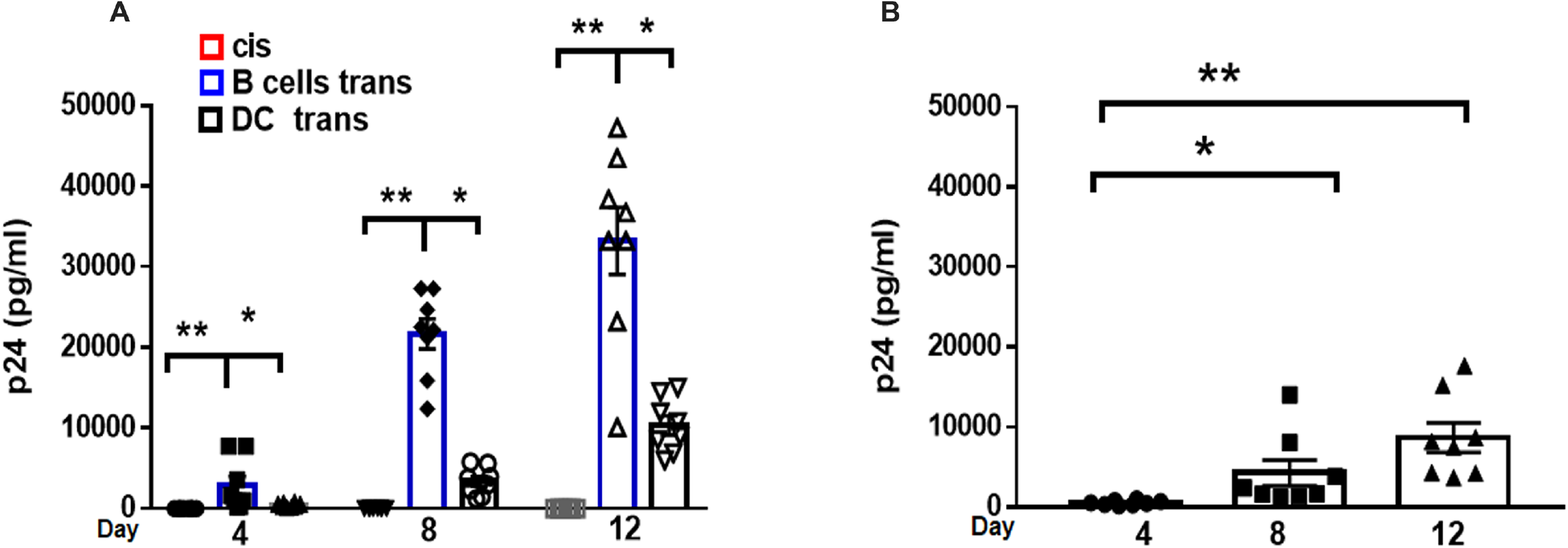
B cells *trans* infect CD4^+^ T cells with higher efficiency than DC. **A.** B lymphocytes and DC were loaded with HIV-1^BaL^ (10^−3^ m.o.i.) as described in Materials and Methods and mixed with PHA/IL2 activated autologous CD4^+^ T cells at a 1:10 ratio for up to 12 days (*trans* infection). CD4^+^ T cells were also pulsed with HIV-1^BaL^ (10^−3^ m.o.i.) and cultured alone (*cis* infection). Coculture supernatants were tested at the indicated time points for HIV Gag p24 levels by ELISA. **B**. CD4^+^ T cells were loaded with HIV-1^BaL^ (10^−1^ m.o.i.) and cultured up to 12 days. Cultures supernatants were tested at the indicated time points for HIV Gag p24 levels by ELISA. Data are mean value ±SE; N=8.; *p<0.05 **p< 0.001.

### B cells, but not DC, *trans* infect naïve CD4^+^ T cells *in vitro*

CD4^+^ T_N_ cells do not express CCR5; however, *in vivo*, T_N_ cells harbor R5-tropic HIV-1 (10, 11, 25, 26). We therefore hypothesized that T_N_ are infected through an APC-mediated *trans* infection mechanism that does not require CCR5 expression by the T cells. To test this hypothesis, we used purified T_N_ and T_CM_ cells as targets for *trans* infection mediated by autologous B lymphocytes or DC that were loaded with 10^−3^ m.o.i. of R5 tropic HIV-1_BaL_. Consistent with the approach described in Figure 1, we initially used PHA/IL2-activated CD4^+^ T_N_ and T_CM_ cells as targets. As shown in Figure 2A, B cells were able to productively *trans* infect either T_N_ or T_CM_ with R5 tropic HIV-1, whereas DC only *trans* infected the T_CM_ subset.

**Figure 2.**
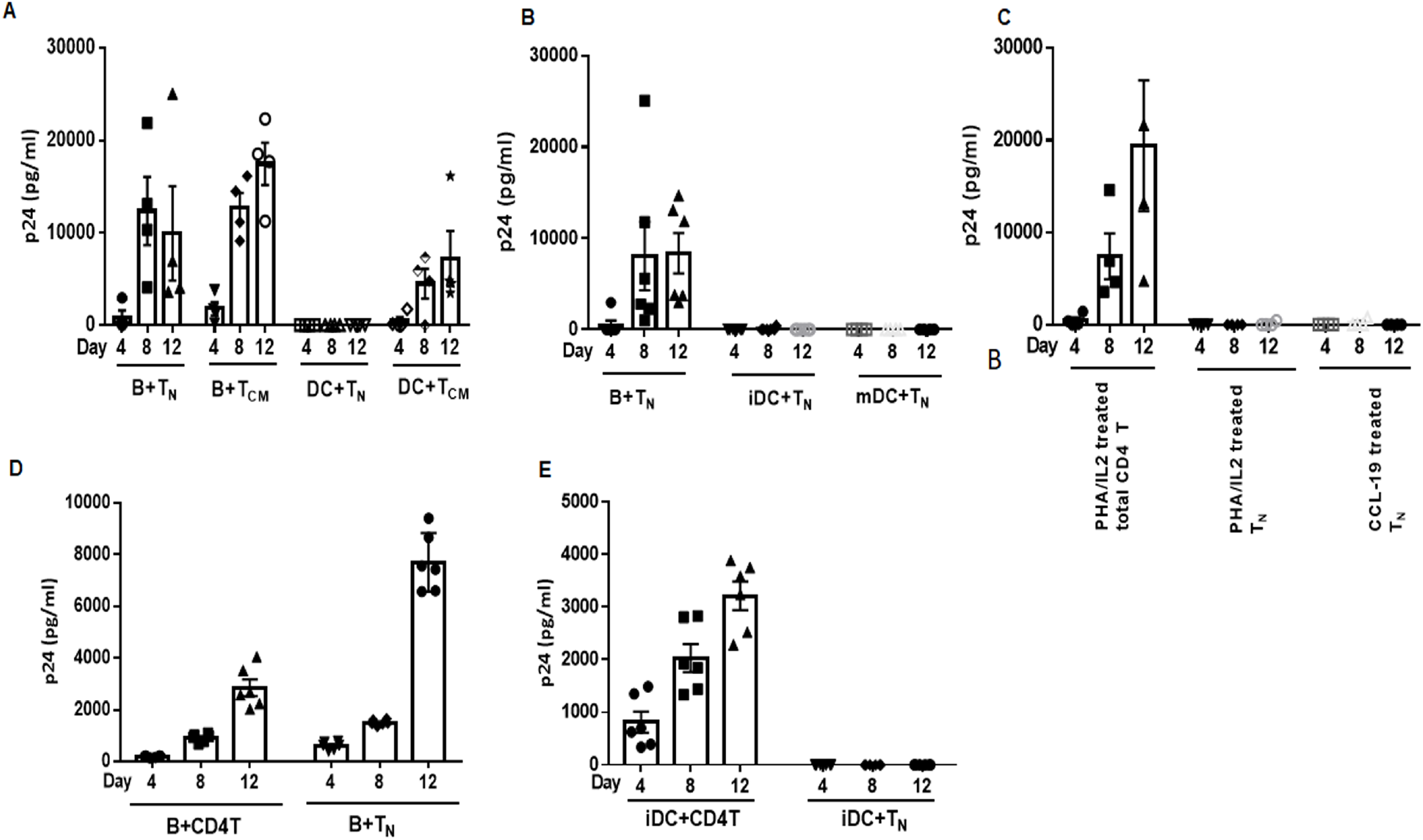
Only B cells *trans* infect T_N_. **A:** B cells or DC were pulsed with HIV-1^BaL^ (10^−3^ m.o.i.) for 2h at 37C, extensively washed and cocultured with PHA/IL2 activated purified naïve (T_N;_ N=5) or central memory (T_CM;_ N=4) CD4^+^T cells. Culture supernatants were collected at the times indicated and tested for HIV Gag p24. Mean±SE. **B.** T_N_ cells were treated with CCL-19, washed and mixed with B cells, immature DC (iDC), or CD40L matured DC (mDC), pulsed with HIV-1^BaL^ (10^−3^ m.o.i.). Cell culture supernatants were collected at the indicated time points and tested for HIV Gag p24 by ELISA. Mean±SE, N=6. **C**: Total CD4^+^ T or T_N_ cells were treated with PHA/IL2 or CCL-19 alone as described in Materials and Methods, washed and directly infected with HIV-1_BAL_ (10^−1^ m.o.i.). Cell culture supernatants were collected at the indicated time points and tested for HIV-1 Gag p24 by ELISA. Mean±SE, N=4 **D.** B cells pulsed with HIV-1 92FR_BX08 (10^−3^ m.o.i.) were mixed with total CD4^+^ or T_N_ cells treated with CCL-19, washed and mixed with as described in Materials and Methods and cultured for up to 12 days. **E.** iDC pulsed with HIV-1 92FR_BX08 (10^−3^ m.o.i.) were mixed with total CD4^+^ or T_N_ cells as described in Materials and Methods and cultured for up to 12 days. Cell cultures supernatants were collected at the indicated time points and tested for HIV-1 Gag p24 by ELISA. Mean±SE, N=6 independent cultures.

PHA/IL2 treatment of CD4^+^ T_N_ and T_CM_ induces T cell activation, thus rendering them more susceptible to HIV-1 infection. Therefore, we next assessed B cell- and DC-mediated HIV-1_BaL_ *trans* infection of T_N_ and T_CM_ cells treated with the chemokine CCL19. As described previously (12), CCL19 neither elicits T cell activation nor induces CCR5 or CXCR4 expression, but enhances *cis* HIV-1 infection of resting CD4^+^ T cells. As shown in Figure 2B, only B cells were able to *trans* infect CCL19-treated T_N_, resulting in detectable HIV-1 Gag p24 in the coculture supernatants. In contrast, neither mDC nor iDC mediated *trans* infection, showing that the ability of DC to *trans* infect T_N_ did not depend on their maturation status. As expected, T_N_ were refractory to direct *cis* infection of HIV-1_BaL_ using either PHA/IL2 or CCL19 conditioned media, while only total CD4^+^ T cells were susceptible to productive *cis* infection (Figure 2C). Our findings were further confirmed using an R5-tropic patient isolate, HIV-1 BX08(92FR_BX08), obtained from the NIAID AIDS Reagent Repository (21). As shown in panels 2D and 2E, B cells could efficiently *trans* infect both total CD4^+^ T and T_N_ cells, while iDC could only *trans* infect total CD4^+^ T cells. *Cis* infection of T_N_ cells was undetectable (not shown). Taken together, these results support that B lymphocytes have a unique ability to mediate highly productive *trans* infection of naïve CD4^+^ T cells with R5-tropic HIV-1.

### Coculture with B cells or DC does not affect the T_N_ phenotype

To address whether the CD4^+^ T_N_ phenotype was altered through coculture with the APC, potentially affecting their efficiency of being *trans* infected with HIV-1, we analyzed T_N_ cells for CCR5 and CD27 expression. T_N_ cultured alone served as a control. CD27 expression was chosen instead of CCR7 expression because CCL19 can induce downregulation of CCR7. The flow cytometry gating strategy is shown in Supplemental Figure S1. As shown in Figure 3A, neither B cells nor DC induced a significantly higher expression of CCR5, up to 12 days in culture (1-way ANOVA), although we detected a slight increase of CCR5 between day 8 and 12 in the DC-T_N_ cocultures. Expression of the CD27 marker also remained unchanged throughout the coculture period (Figure 3B).

**Figure 3.**
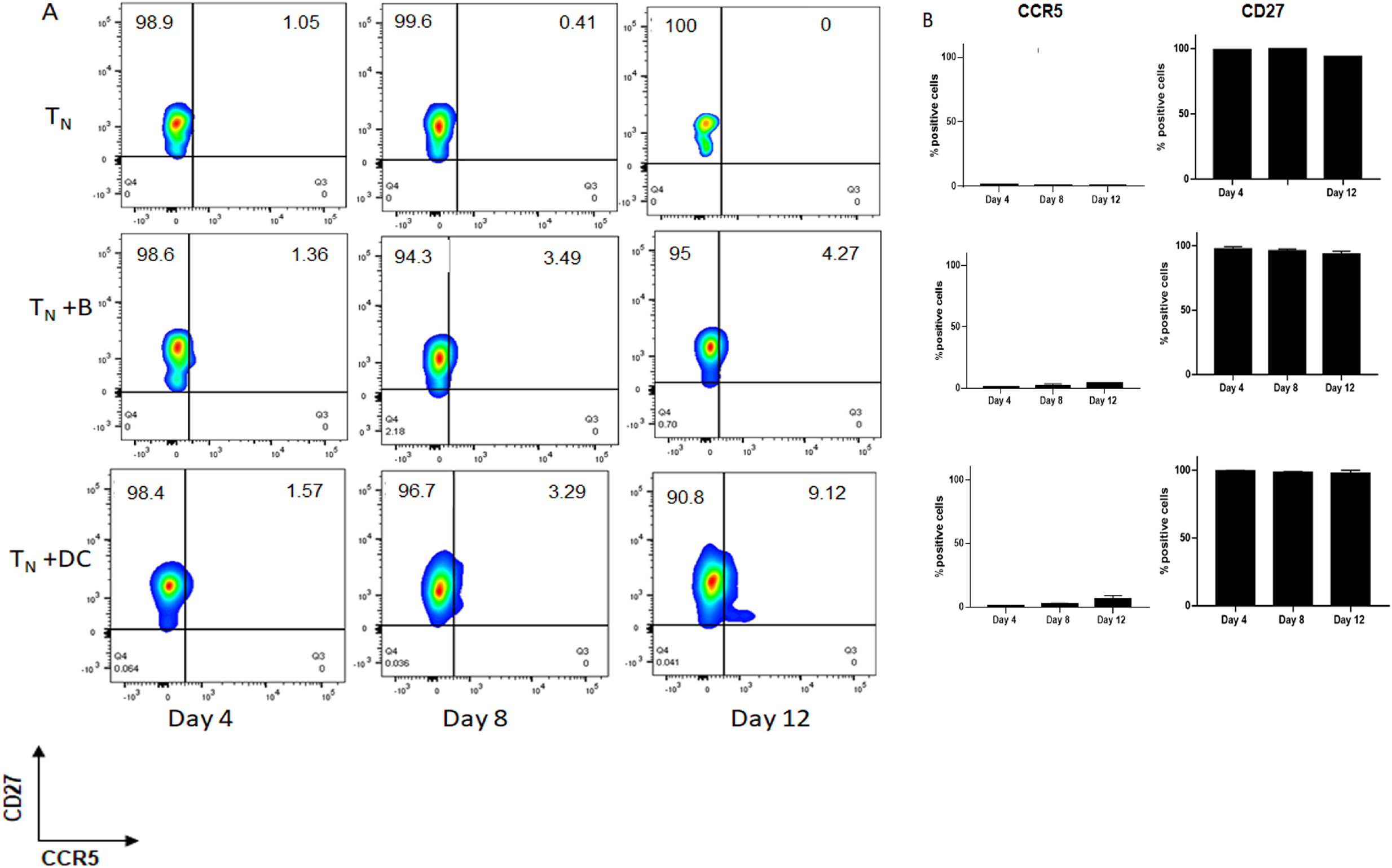
Coculture with B cells or DC does not affect T_N_ phenotype. **A.** T_N_ cells were cultured alone (top row) or cocultured with HIV-1^BaL^ pulsed B cells or DC (middle and bottom row, respectively), sampled at the indicated time points, stained with anti-CCR5 and CD27 mAB and analyzed by FACS as described in Materials and Methods. Representative data from 3 independent experiments. **B.** CCR5 and CD27 percent positive cells in T_N_ control culture or cocultures. Mean±SE, N=3.

These data show that there is no significant alteration of the T_N_ phenotype during coculture with APCs, and confirm that B cells can establish HIV-1_BaL_ infection in T_N_ in the absence of significant CCR5 coreceptor expression.

### Detection of intracellular HIV-1 p24 in APC-T_N_ cocultures

Given the slight increase of CCR5 in the DC-T_N_ cocultures, we next questioned if the detection of HIV-1 p24 we measured in the *trans* infection coculture supernatant reflected p24 intracellular localization. We therefore stained cells collected from the *trans* infection wells and examined them for p24 expression by flow cytometry. As shown in Figure 4, we were able to detect intracellular HIV-1 p24 in the cocultures of B cells with either total CD4^+^ T or T_N_. In the DC mediated *trans* infection cocultures, we could only detect HIV-1 p24 in the DC-total CD4^+^T cell wells, with very low levels in the DC-T_N_ cocultures. Taken together, these data further support the conclusion that only B lymphocytes can efficiently *trans* infect T_N_ cells.

**Figure 4.**
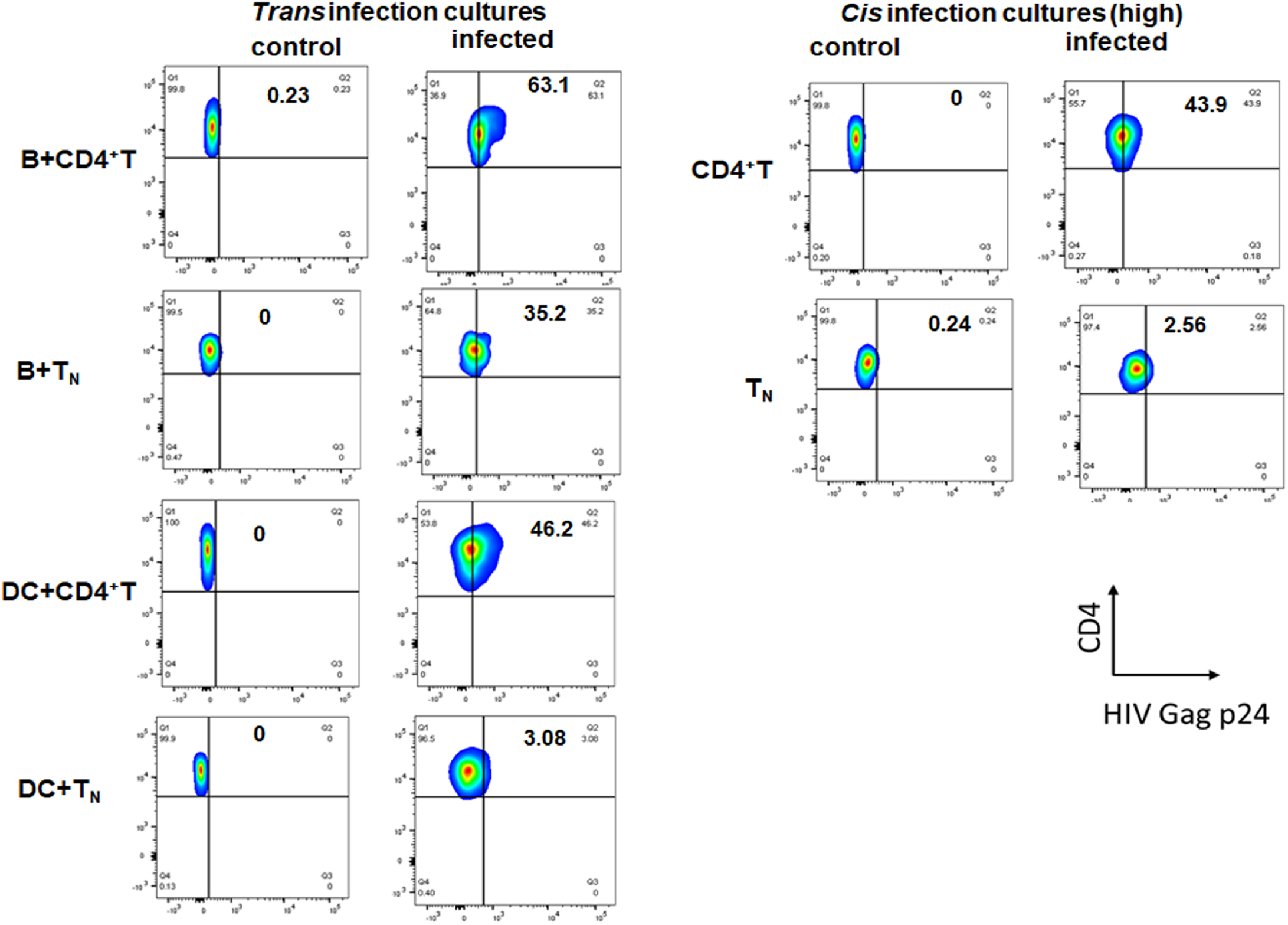
Detection of HIV-1 p24 antigen in *trans* infection coculture with B cells or DC: B cells or DC loaded with HIV-1 (10^−3^ m.o.i) were cultured with total CD4^+^ T or T_N_ cells, sampled after 8 days, and stained with anti-Kc57, -CD4, -CD3, or -CD19 and analyzed by FACS as described in Materials and Methods.. ***Cis* infection:** CD4^+^ T or T_N_ cells were infected with HIV-1^BaL^(10^−1^ m.o.i.) cultured, stained, and analyzed by flow cytometry in parallel to the *trans* infection cocultures. Representative data from 2 independent experiments.

### B cell *trans* infection of T_N_ cells is mediated by DC-SIGN

We have previously shown that B cell *trans* infection of CD4^+^T cells is inhibited by blocking of DC-SIGN (15). Therefore, we tested if DC-SIGN was also necessary to mediated *trans* infection of T_N_ cells. As shown in Figure 5A, blocking of B cells with anti-DC-SIGN mAb significantly inhibited *trans* infection of T_N_. We also treated T_N_ cells with maraviroc, a chemokine co-receptor 5 (CCR5) antagonist, to determine if any amount of CCR5 expressed by T_N_ in the trans infection cocultures could be responsible for the infection. As expected, treatment with maraviroc did not significantly inhibit the efficient *trans* infection of T_N_ (Figure 5A). These data were also confirmed by HIV-1 p24 intracellular staining of *trans* infection cocultures (Figure 5B).

**Figure 5.**
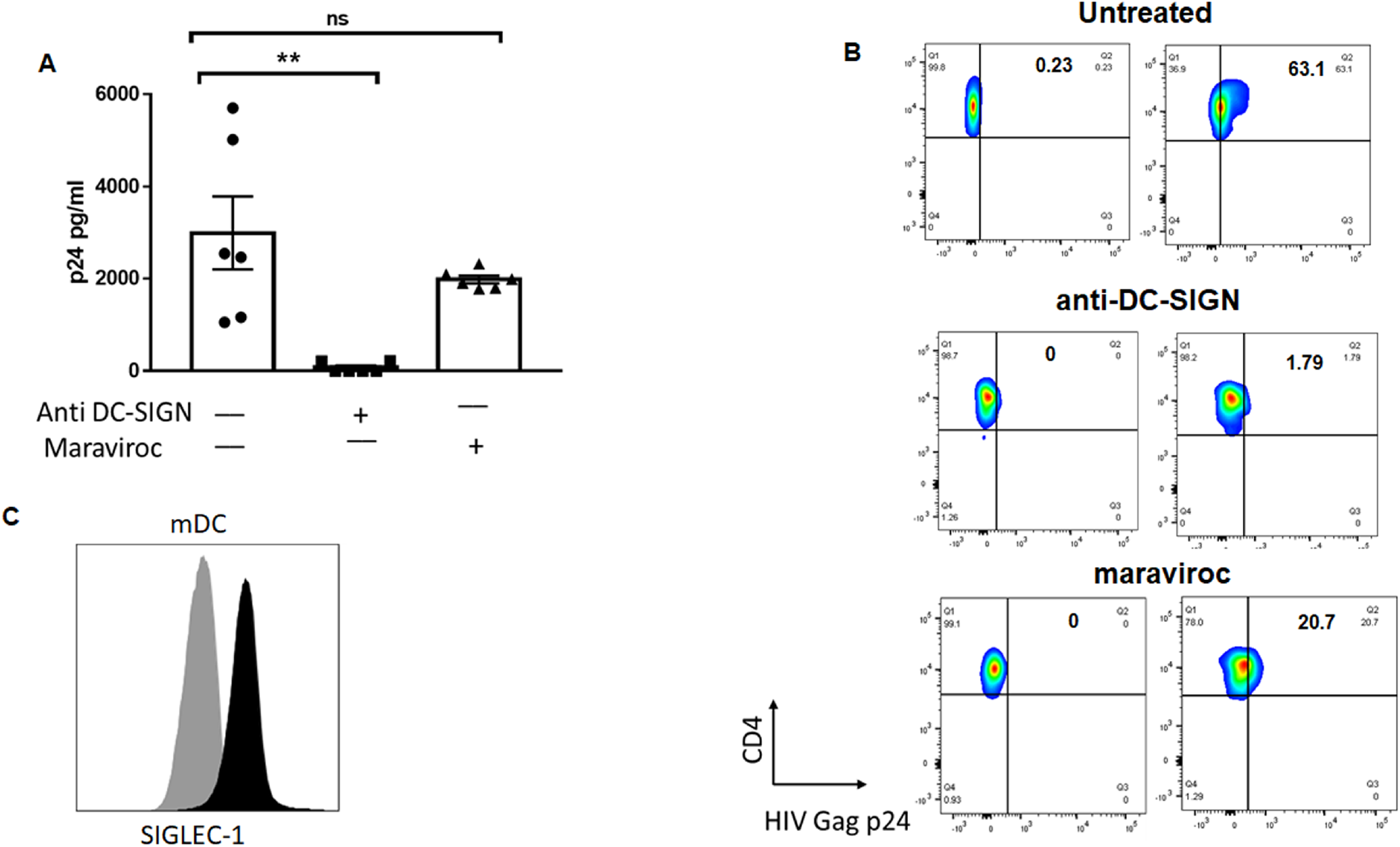
B cell-mediated *trans* infection of T_N_ cells is inhibited by anti-DC-SIGN blocking. **A.** B cells were incubated with 20μg/ml anti DC-SIGN mAb for 1h at 4C prior to pulsing with HIV-1^BaL^ (10^−3^ m.o.i.) and cocultured with T_N_ cells for *trans* infection as described in Materials and Methods. Supernatants were collected after 12 days and tested for HIV Gag p24 by ELISA. B cells treated with mouse IgG (20μg/ml) were used as untreated control. Mean±SE, N=6 independent cultures. **B.** In parallel cultures, T_N_ cells were treated with Maraviroc (1uM) as described in Materials and Methods and cocultured with HIV-1^BaL^ pulsed B cells for *trans* infection. Supernatants were collected after 12 days and tested for HIV-1 Gag p24 by ELISA. Mean±SE, N=6 independent cultures. **C.** DC matured with CD40L/IL4 were collected and stained for CD169 (Siglec-1) expression (black histogram) and compared to an isotype control (grey histogram) isotype control. Representative of 2 independent experiments.

It has been proposed that siglec-1 (CD169) is a key factor for efficient DC-mediated HIV-1 *trans* infection with DC matured by lipopolysaccharide (28). Since our DC were used as immature cells or matured with CD40L/IL4, we tested if the inefficient *trans* infection of T_N_ cells by DC was due to lack of siglec-1 expression, although both B cells and DC could efficiently *trans* infect total CD4^+^ T cells. We found that CD40L/IL4-matured DC expressed siglec-1 (Figure 5C), excluding the possibility that that this receptor contributed to the observed phenotype.

### Reactivation of HIV-1 from T_N_

To assess whether HIV-1 *trans* infection of T_N_ cells mediated by B cells or DC resulted in HIV-1 latency, T_N_ cells were cultured with either HIV-1_BaL_-loaded B cells or DC for 8 days and then treated with PMA/PHA, and culture supernatants were harvested every 3 days for p24 analysis (Fig.6A). As shown in Figure 6B, HIV-1 was recovered only from the B-T_N_ cocultures. This indicates that the lack of detectable virus replication in the DC-T_N_ was not due to the establishment of latency without detectable viral replication in the T_N_.

**Figure 6.**
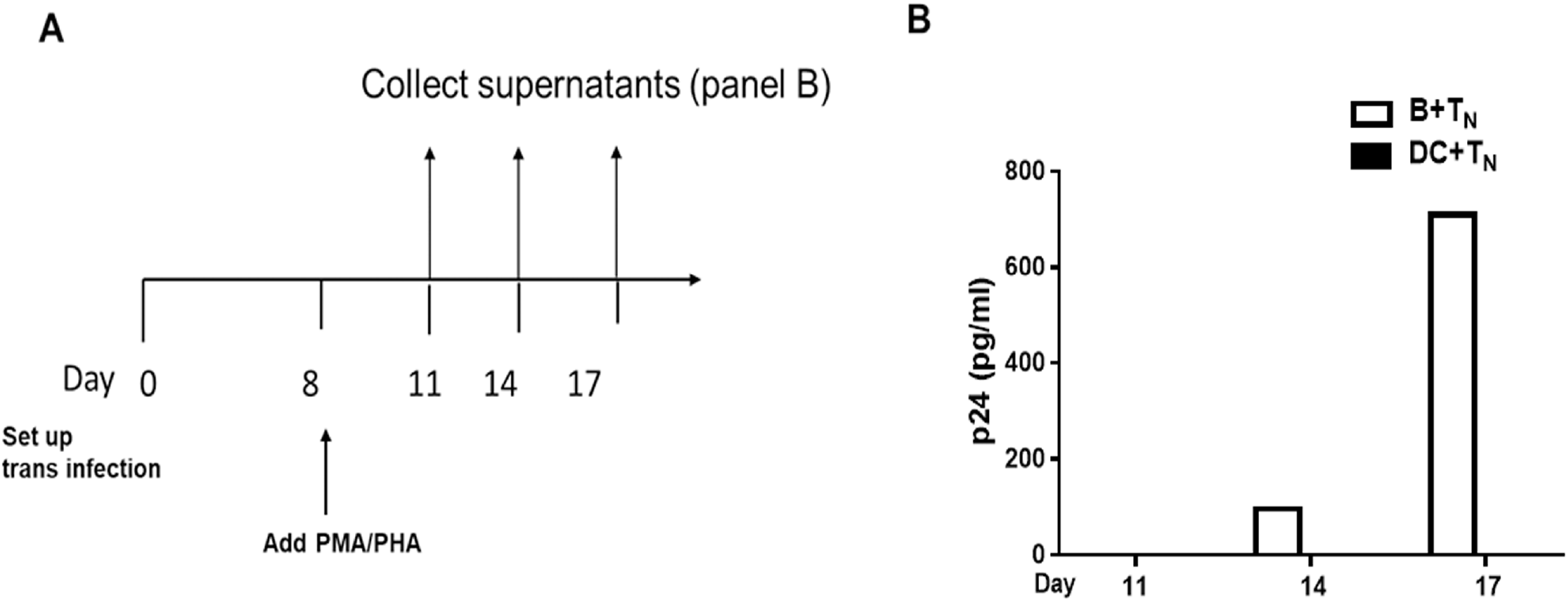
Detection of virus after LRA reactivation. **A.** Schematic representation of the experimental approach to measure reversal of HIV-1 latency in T_N_ cells *trans* infected by B cells or DC. **B.** cocultures treated with LRA activators at day 8 and then sampled at the indicated time point after reactivation. Supernatants were tested for HIV Gag p24 by ELISA.

### CD4^+^ T_N_ cells from HIV-1-infected NP harbor less total HIV-1 DNA

We have previously shown that APC from NP cannot *trans* infect autologous and heterologous CD4^+^ T cells, and that this phenotype is under control of cellular cholesterol homeostasis regulation (18, 19). Furthermore, this characteristic is present prior to infection with HIV-1, indicating that it is an innate, genetically controlled phenotype. If B cell-mediated *trans* infection of T_N_ is an important mechanism by which these cells become infected with HIV-1, then it is plausible that NP have a reduced or absent level of HIV-1 DNA in this CD4^+^ T cell subset. We therefore quantified the viral DNA reservoir in total CD4^+^ T cells and T_N_ cells from 7 NP not under ART at the time of testing, and 7 PR on ART (Figure 7). The results show that we could not detect HIV-1 DNA in T_N_ from NP classified as elite controllers (EC), while a relatively low number of HIV-1 DNA copies were detected in LTNP and VC (19). Overall, the average copy number of HIV-1 DNA in T_N_ from NP was lower compared to the number of copies detected in the 7 PR ART-suppressed participants (p = 0.007). Both NP and PR had similar levels of HIV-1 DNA copies when total CD4^+^ T cells were tested. Given that CD4^+^ T cells from NPs are susceptible to direct, *cis* infection as well as CD4^+^T cells from PR (19), the evidence supports the concept that the low amount of HIV-1 DNA detected is the result of direct infection. Taken together, these data suggest that individuals naturally able to control HIV-1 disease progression have a reduced or absent HIV-1 reservoir in their T_N_ population.

**Figure7.**
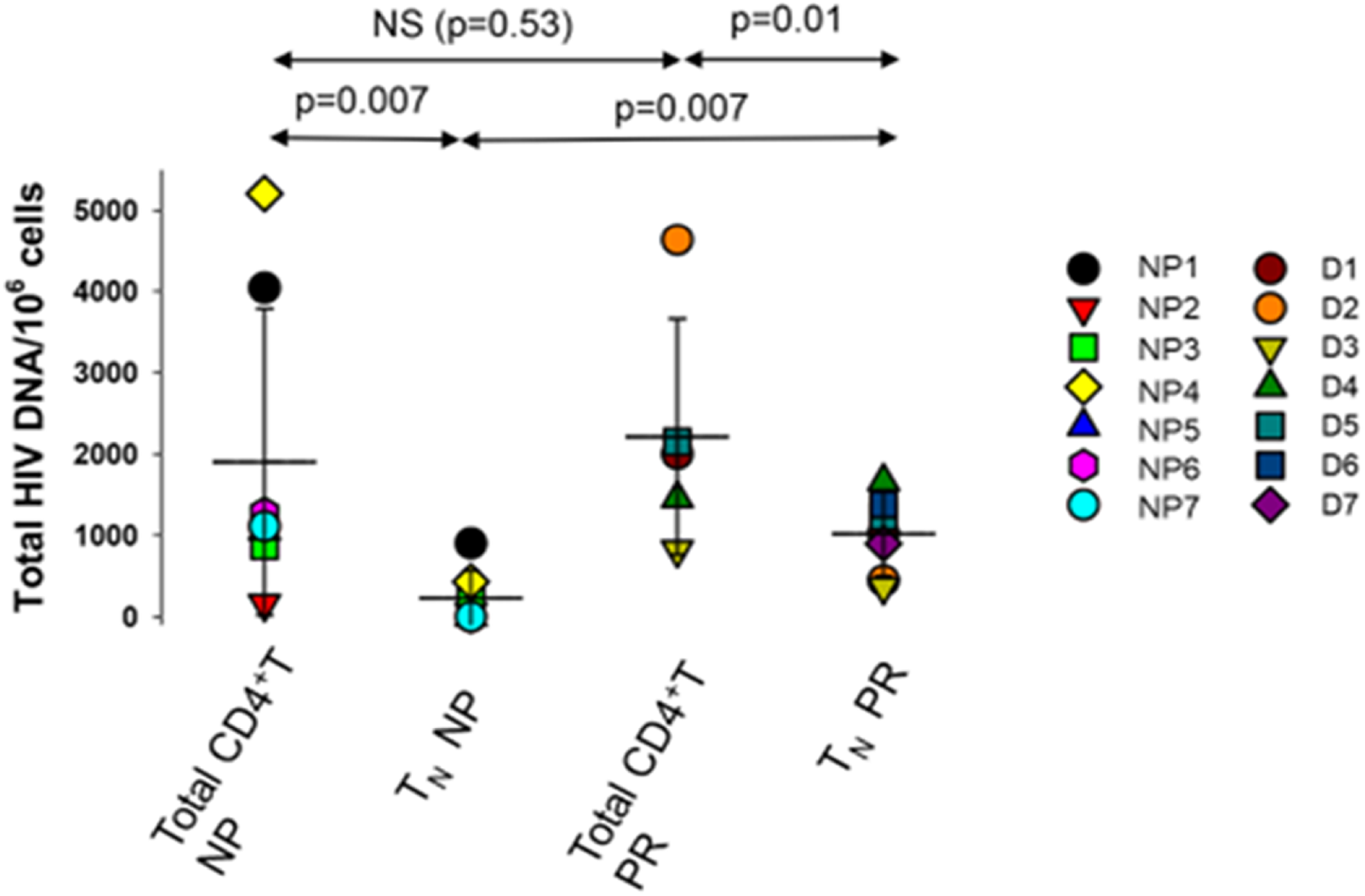
Quantification of total HIV-1 DNA in CD4^+^ total and T_N_ cells. Each dot represents a unique donor. Statistical comparison was analyzed using a Wilcoxon matched-pairs signed rank test. A p value < 0.05 was considered significant

## Discussion

Here we show that B lymphocytes have the unique ability to *trans* infect CD4^+^ T_N_ cells *in vitro* with an R5 tropic HIV-1 laboratory strain (HIV-1_BaL_) and an R5 clinical isolate (HIV-1 BX08(92FR_BX08)), compared to myeloid DC. Prior studies have shown that T_N_ cells can be infected *in vitro* with CXCR4-tropic HIV-1 when pretreated with the chemokine CCL19, the ligand for the CCR7 receptor, which expression significantly increases during the acute phase of infection when the latent reservoir is established (9, 29). This treatment does not alter the activation or proliferation state of T_N_ cells, and does not induce significant expression of the CCR5 co-receptor. Therefore, this model was used in our study to preserve the phenotype of the T_N_ population, which remained resistant to *cis* infection with an R-5 tropic HIV strain. Furthermore, exposure of T_N_ cells to B cells loaded with R-5 tropic HIV-1_BaL_ during the coculture period did not induce higher expression of the CCR5 receptor, thus excluding the possibility that the efficient *trans* infection we observed was the result of *in vitro* conditions. Efficient transfer of HIV-1 to T_N_ mediated by B cells was also confirmed by detection of intracellular HIV-1 p24 by flow cytometry.

We have previously shown that activated B cells are able to bind and internalize HIV-1 into cytoplasmic vesicles through DC-SIGN (15), and can *trans* infect CD4^+^ T cells for up to two days with high efficiency. Furthermore, we have demonstrated that *trans* infection of total CD4^+^ T cells can be inhibited by treatment with anti-DC-SIGN mAb. Here, we have confirmed that inhibition of DC-SIGN expression on B cells also impairs *trans* infection of T_N_. Notably, B cells do not support HIV-1 replication (15). Therefore, the second, *cis* infection phase in DC-mediated HIV-1 *trans* infection (30) is not applicable to B lymphocytes. Since cell-to-cell mediated spread of HIV-1 is several orders of magnitude more efficient compared to direct *cis* infection of target cells (17), this mode of virus dissemination could have a significant role in HIV pathogenesis, particularly in T cell-APC dense anatomical compartments (16, 31). We propose that this *trans* infection process is likely intertwined with basic immunologic interactions of B lymphocytes and naïve T cells. Indeed, B cells were recently described as having a broad role in the development of T_N_ cells (32). The interaction between B cells and T_N_ cells thus goes beyond the classical initiation of antigen-specific B cell differentiation into antibody producing plasma cells. In fact, evidence suggests that B cells are necessary and sufficient to prime and activate T_N_ cells in response to virus-like particles (32). Thus, unique features of interactions between B and T_N_ cells could drive the transfer and replication of HIV-1. Notably, it is known that B cells are superior to DC in capturing high doses of cognate antigen through high affinity antigen-specific receptors, therefore rendering B cell-mediated antigen stimulation more efficient than DC (33). In T-dependent B cell immune responses, antigen-engaged B cells must find their cognate helper T cells to initiate the progression of B cell immune responses. Within the lymph node follicle, B cells move continuously to survey the subcapsular (SCS) macrophages for surface-displayed antigens (34, 35) and are also receiving survival signals from fibroblastic reticular cells (FRC) (36) such as the B cell activator BAFF, which has been shown to activate B cells to express DC-SIGN (37). These B cells are positioned to capture HIV-1 either as free virus entering through the afferent lymph vessel or through sampling of SCS macrophages (33) which have poor endocytic capacity and limited degradative ability (35). This ultimately prevents them from efficiently degrading HIV-1. In this scenario, subcapsular B cells activated through a T-independent mechanism are perfectly positioned to capture HIV-1 particles while surveying the environment for their specific antigen.

Upon encounter with antigen, signaling via the B cell receptor (BCR) starts the sequence of events that will bring the antigen-specific B cells to the follicle-T zone boundary where they will search for their cognate CD4^+^ T cell among the T_N_ cells residing there (38). This interaction provides an opportunity for transfer of HIV-1 that has been captured by B cells to the CD4^+^ T cells This interaction at the follicle-T cell zone interface of lymph nodes can last from several minutes to an hour (39, 40), and requires the interaction of integrins, such as LFA1 on T helper cells interacting with ICAM-1 or ICAM-2 on B cells, as well as co-stimulatory molecule CD86 signaling of CD28. Crucially, these interactions are stabilized by the antigenic peptide presented by B cell-expressed MHC class II. B cells capture antigen with high affinity through the B cell receptor (BCR), allowing for even low concentration of antigen to result in high internalization and subsequent presentation to T cells (41) and upregulation of the costimulatory molecule CD86 expression. On the other hand, DC capture antigen through nonspecific binding, requiring higher levels of antigen to induce a CD4^+^ T cell response. Moreover, the interaction between DC and T_N_ helper cells is not as long lived, resulting in a lower chance of virus being transferred, even though activated B cell and DC express costimulatory molecules involved in the formation of the immunological synapse. Thus, the unique features of the interactions between B and T_N_ cells could drive the transfer of HIV-1 to T_N_ with higher efficiency compared to DC.

Although T_N_ cells represent the more abundant fraction of CD4^+^ T cells, most studies of the latent HIV-1 reservoir have focused on T_M_ cells because they harbor the highest levels of HIV-1 DNA in people under ART. We and others (9, 12, 26) have shown that although the frequency of HIV-1 infection in these cells is lower compared to other subsets, as much or more virus is produced by these cells after treatment with latency-reactivation agents (LRA). This is true also when T_N_ cells isolated from HIV-1 infected individuals under ART are exposed to LRA. Paradoxically, although T_N_ do not express the HIV-1 co-receptor CCR5, they harbor CCR5-tropic virus *in vivo*. Therefore, an understanding on how this subset of CD4^+^ T cells becomes infected could provide important clues in the development of strategies to thwart the early establishment of HIV-1 infection.

As we have previously shown, efficient APC-mediated *trans* infection is regulated by APC membrane cholesterol content, and is related to the control of HIV-1 disease progression (18, 19). In fact, APC derived from HIV-1-infected NP have an innate inability to *trans* infect CD4^+^ T cells, and this phenotype can be reversed by replenishing cell membrane cholesterol. On the other hand, APC from HIV-1-infected individuals with progressing disease, i.e., PR, mediate efficient HIV-1 *trans* infection (18, 19). Here we quantified the viral reservoirs in total CD4^+^ T and T_N_ from NP and PR in the Pittsburgh clinical site of the MWCCS. Notably, while the PR studied here were under suppressive ART, all the NP tested were therapy naïve at the time of the study. We could not detect viral DNA in NPs classified as EC, while a significantly smaller number of HIV-1 DNA copies was detected in LTNP and VC compared to PR.These data strongly suggest that the altered ability to *trans* infect CD4^+^ T cells in NP results in a small or negligible pool of latently infected T_N_ cells, thus contributing to the maintenance of the NP phenotype, at levels even lower than those detectable in patients under suppressive ART. Although limited in scope, our findings are also consistent with that observed in the French Virological and Immunological Studies in Controllers After Treatment Interruption (VISCONTI) cohort of individuals that received ART within 10 weeks of primary infection (42), where viremia was controlled for 24 months post-treatment interruption. In that cohort, HIV-1 DNA was detected in T_N_ of only 2 out of 11 patients, while the other T cell subsets harbored comparable levels of HIV-1 DNA. Our present study suggests that early, B cell-mediated *trans* infection could be an important mechanism by which HIV-1, regardless of its basic cell tropism, establishes infection in T_N_ cells. We propose an additional role for B cell-mediated *trans* infection, not only as an efficient means to spread HIV-1 to CD4^+^ T cells, but as the driver in establishing the HIV-1 reservoir in T_N_ and potential consequent control of HIV-1 disease progression.

## Acknowledgments

The authors swish to thank Kathleen Hartle and Patrick Mehta for technical assistance and the Pitt Men’s Study staff and volunteers for their support. This work was supported by the National Institute of Health grants R01AI118403, U01-AI35041, U01 HL146208, T32AI065380.

---

**Supplemental Figure 1. Gating Strategy to determine T_N_ phenotype** Lymphocytes were gated first based on forward and side scatter, followed by doublet event exclusion, then by exclusion of dead cells (Aqua dye positive). CD4^+^ positive cells were then gated into CD45RA negative and CD45RA positive populations, with the latter population being 100% CCR7 positive as well as CCR5 negative.

**Supplemental Figure 2. Gating Strategy to determine intracellular HIV-1 p24.** Lymphocytes were gated first based on forward and side scatter, followed by doublet event exclusion, then by exclusion of dead cells (Aqua dye positive). CD4^+^/HIV p24^+^ cells were then gated within the CD4^+^/CD3^+^ population

## References

1. Chun TW, Stuyver L, Mizell SB, Ehler LA, Mican JA, Baseler M, Lloyd AL, Nowak MA, Fauci AS. 1997. Presence of an inducible HIV-1 latent reservoir during highly active antiretroviral therapy. Proc Natl Acad Sci U S A 94:13193–7.

2. Eisele E, Siliciano RF. 2012. Redefining the viral reservoirs that prevent HIV-1 eradication. Immunity 37:377–88.

3. Finzi D, Blankson J, Siliciano JD, Margolick JB, Chadwick K, Pierson T, Smith K, Lisziewicz J, Lori F, Flexner C, Quinn TC, Chaisson RE, Rosenberg E, Walker B, Gange S, Gallant J, Siliciano RF. 1999. Latent infection of CD4+ T cells provides a mechanism for lifelong persistence of HIV-1, even in patients on effective combination therapy. Nat Med 5:512–7.

4. Finzi D, Hermankova M, Pierson T, Carruth LM, Buck C, Chaisson RE, Quinn TC, Chadwick K, Margolick J, Brookmeyer R, Gallant J, Markowitz M, Ho DD, Richman DD, Siliciano RF. 1997. Identification of a reservoir for HIV-1 in patients on highly active antiretroviral therapy. Science 278:1295–300.

5. Chomont N, El-Far M, Ancuta P, Trautmann L, Procopio FA, Yassine-Diab B, Boucher G, Boulassel MR, Ghattas G, Brenchley JM, Schacker TW, Hill BJ, Douek DC, Routy JP, Haddad EK, Sekaly RP. 2009. HIV reservoir size and persistence are driven by T cell survival and homeostatic proliferation. Nat Med 15:893–900.

6. Schnittman SM, Lane HC, Greenhouse J, Justement JS, Baseler M, Fauci AS. 1990. Preferential infection of CD4+ memory T cells by human immunodeficiency virus type 1: evidence for a role in the selective T-cell functional defects observed in infected individuals. Proc Natl Acad Sci U S A 87:6058–62.

7. Dai J, Agosto LM, Baytop C, Yu JJ, Pace MJ, Liszewski MK, O’Doherty U. 2009. Human immunodeficiency virus integrates directly into naive resting CD4+ T cells but enters naive cells less efficiently than memory cells. J Virol 83:4528–37.

8. Zerbato JM, Purves HV, Lewin SR, Rasmussen TA. 2019. Between a shock and a hard place: challenges and developments in HIV latency reversal. Curr Opin Virol 38:1–9.

9. Zerbato JM, Serrao E, Lenzi G, Kim B, Ambrose Z, Watkins SC, Engelman AN, Sluis-Cremer N. 2016. Establishment and Reversal of HIV-1 Latency in Naive and Central Memory CD4+ T Cells In Vitro. J Virol 90:8059–73.

10. Ostrowski MA, Chun TW, Justement SJ, Motola I, Spinelli MA, Adelsberger J, Ehler LA, Mizell SB, Hallahan CW, Fauci AS. 1999. Both memory and CD45RA+/CD62L+ naive CD4(+) T cells are infected in human immunodeficiency virus type 1-infected individuals. J Virol 73:6430–5.

11. Wightman F, Solomon A, Khoury G, Green JA, Gray L, Gorry PR, Ho YS, Saksena NK, Hoy J, Crowe SM, Cameron PU, Lewin SR. 2010. Both CD31(+) and CD31(-) naive CD4(+) T cells are persistent HIV type 1-infected reservoirs in individuals receiving antiretroviral therapy. J Infect Dis 202:1738–48.

12. Zerbato JM, McMahon DK, Sobolewski MD, Mellors JW, Sluis-Cremer N. 2019. Naive CD4+ T Cells Harbor a Large Inducible Reservoir of Latent, Replication-Competent HIV-1. Clin Infect Dis doi:10.1093/cid/ciz108.

13. Cameron PU, Freudenthal PS, Barker JM, Gezelter S, Inaba K, Steinman RM. 1992. Dendritic cells exposed to human immunodeficiency virus type-1 transmit a vigorous cytopathic infection to CD4+ T cells. Science 257:383–7.

14. Geijtenbeek TB, Kwon DS, Torensma R, van Vliet SJ, van Duijnhoven GC, Middel J, Cornelissen IL, Nottet HS, KewalRamani VN, Littman DR, Figdor CG, van Kooyk Y. 2000. DC-SIGN, a dendritic cell-specific HIV-1-binding protein that enhances trans-infection of T cells. Cell 100:587–97.

15. Rappocciolo G, Piazza P, Fuller CL, Reinhart TA, Watkins SC, Rowe DT, Jais M, Gupta P, Rinaldo CR. 2006. DC-SIGN on B lymphocytes is required for transmission of HIV-1 to T lymphocytes. PLoS Pathog 2:e70.

16. Sigal A, Baltimore D. 2012. As good as it gets? The problem of HIV persistence despite antiretroviral drugs. Cell Host Microbe 12:132–8.

17. Zhong P, Agosto LM, Ilinskaya A, Dorjbal B, Truong R, Derse D, Uchil PD, Heidecker G, Mothes W. 2013. Cell-to-cell transmission can overcome multiple donor and target cell barriers imposed on cell-free HIV. PLoS One 8:e53138.

18. DeLucia DC, Rinaldo CR, Rappocciolo G. 2018. Inefficient HIV-1 trans Infection of CD4(+) T Cells by Macrophages from HIV-1 Nonprogressors Is Associated with Altered Membrane Cholesterol and DC-SIGN. J Virol 92.

19. Rappocciolo G, Jais M, Piazza P, Reinhart TA, Berendam SJ, Garcia-Exposito L, Gupta P, Rinaldo CR. 2014. Alterations in cholesterol metabolism restrict HIV-1 trans infection in nonprogressors. MBio 5:e01031–13.

20. Sankapal S, Gupta P, Ratner D, Ding M, Shen C, Sanyal A, Stolz D, Cu-Uvin S, Ramratnam B, Chen Y. 2016. HIV Exposure to the Epithelia in Ectocervical and Colon Tissues Induces Inflammatory Cytokines Without Tight Junction Disruption. AIDS Res Hum Retroviruses 32:1054–1066.

21. Brown BK, Darden JM, Tovanabutra S, Oblander T, Frost J, Sanders-Buell E, de Souza MS, Birx DL, McCutchan FE, Polonis VR. 2005. Biologic and genetic characterization of a panel of 60 human immunodeficiency virus type 1 isolates, representing clades A, B, C, D, CRF01_AE, and CRF02_AG, for the development and assessment of candidate vaccines. J Virol 79:6089–101.

22. Rappocciolo G, Sluis-Cremer N, Rinaldo CR. 2019. Efficient HIV-1 Trans Infection of CD4(+) T Cells Occurs in the Presence of Antiretroviral Therapy. Open Forum Infect Dis 6:ofz253.

23. Hong F, Aga E, Cillo AR, Yates AL, Besson G, Fyne E, Koontz DL, Jennings C, Zheng L, Mellors JW. 2016. Novel Assays for Measurement of Total Cell-Associated HIV-1 DNA and RNA. J Clin Microbiol 54:902–11.

24. Rappocciolo G, Jenkins FJ, Hensler HR, Piazza P, Jais M, Borowski L, Watkins SC, Rinaldo CR, Jr. 2006. DC-SIGN is a receptor for human herpesvirus 8 on dendritic cells and macrophages. J Immunol 176:1741–9.

25. Heeregrave EJ, Geels MJ, Brenchley JM, Baan E, Ambrozak DR, van der Sluis RM, Bennemeer R, Douek DC, Goudsmit J, Pollakis G, Koup RA, Paxton WA. 2009. Lack of in vivo compartmentalization among HIV-1 infected naive and memory CD4+ T cell subsets. Virology 393:24–32.

26. Venanzi Rullo E, Cannon L, Pinzone MR, Ceccarelli M, Nunnari G, O’Doherty U. 2019. Genetic Evidence That Naive T Cells Can Contribute Significantly to the Human Immunodeficiency Virus Intact Reservoir: Time to Re-evaluate Their Role. Clin Infect Dis 69:2236–2237.

27. Sigal A, Kim JT, Balazs AB, Dekel E, Mayo A, Milo R, Baltimore D. 2011. Cell-to-cell spread of HIV permits ongoing replication despite antiretroviral therapy. Nature 477:95–8.

28. Izquierdo-Useros N, Lorizate M, Puertas MC, Rodriguez-Plata MT, Zangger N, Erikson E, Pino M, Erkizia I, Glass B, Clotet B, Keppler OT, Telenti A, Krausslich HG, Martinez-Picado J. 2012. Siglec-1 is a novel dendritic cell receptor that mediates HIV-1 trans-infection through recognition of viral membrane gangliosides. PLoS Biol 10:e1001448.

29. Saleh S, Solomon A, Wightman F, Xhilaga M, Cameron PU, Lewin SR. 2007. CCR7 ligands CCL19 and CCL21 increase permissiveness of resting memory CD4+ T cells to HIV-1 infection: a novel model of HIV-1 latency. Blood 110:4161–4.

30. Turville SG, Santos JJ, Frank I, Cameron PU, Wilkinson J, Miranda-Saksena M, Dable J, Stossel H, Romani N, Piatak M, Jr., Lifson JD, Pope M, Cunningham AL. 2004. Immunodeficiency virus uptake, turnover, and 2-phase transfer in human dendritic cells. Blood 103:2170–9.

31. Sattentau Q. 2008. Avoiding the void: cell-to-cell spread of human viruses. Nat Rev Microbiol 6:815–26.

32. Hong S, Zhang Z, Liu H, Tian M, Zhu X, Zhang Z, Wang W, Zhou X, Zhang F, Ge Q, Zhu B, Tang H, Hua Z, Hou B. 2018. B Cells Are the Dominant Antigen-Presenting Cells that Activate Naive CD4(+) T Cells upon Immunization with a Virus-Derived Nanoparticle Antigen. Immunity 49:695–708 e4.

33. Cyster JG. 2010. B cell follicles and antigen encounters of the third kind. Nat Immunol 11:989–96.

34. Batista FD, Harwood NE. 2009. The who, how and where of antigen presentation to B cells. Nat Rev Immunol 9:15–27.

35. Carrasco YR, Batista FD. 2007. B cells acquire particulate antigen in a macrophage-rich area at the boundary between the follicle and the subcapsular sinus of the lymph node. Immunity 27:160–71.

36. Katakai T, Suto H, Sugai M, Gonda H, Togawa A, Suematsu S, Ebisuno Y, Katagiri K, Kinashi T, Shimizu A. 2008. Organizer-like reticular stromal cell layer common to adult secondary lymphoid organs. J Immunol 181:6189–200.

37. He B, Qiao X, Klasse PJ, Chiu A, Chadburn A, Knowles DM, Moore JP, Cerutti A. 2006. HIV-1 envelope triggers polyclonal Ig class switch recombination through a CD40-independent mechanism involving BAFF and C-type lectin receptors. J Immunol 176:3931–41.

38. Cyster JG, Allen CDC. 2019. B Cell Responses: Cell Interaction Dynamics and Decisions. Cell 177:524–540.

39. Allen CD, Okada T, Cyster JG. 2007. Germinal-center organization and cellular dynamics. Immunity 27:190–202.

40. Qi H, Kastenmuller W, Germain RN. 2014. Spatiotemporal basis of innate and adaptive immunity in secondary lymphoid tissue. Annu Rev Cell Dev Biol 30:141–67.

41. Zaretsky I, Atrakchi O, Mazor RD, Stoler-Barak L, Biram A, Feigelson SW, Gitlin AD, Engelhardt B, Shulman Z. 2017. ICAMs support B cell interactions with T follicular helper cells and promote clonal selection. J Exp Med 214:3435–3448.

42. Saez-Cirion A, Bacchus C, Hocqueloux L, Avettand-Fenoel V, Girault I, Lecuroux C, Potard V, Versmisse P, Melard A, Prazuck T, Descours B, Guergnon J, Viard JP, Boufassa F, Lambotte O, Goujard C, Meyer L, Costagliola D, Venet A, Pancino G, Autran B, Rouzioux C, Group AVS. 2013. Post-treatment HIV-1 controllers with a long-term virological remission after the interruption of early initiated antiretroviral therapy ANRS VISCONTI Study. PLoS Pathog 9:e1003211.

